# DNA Cut-Ligation Cyclization Surpasses Jacobson-Stockmayer J-Factor Expectations by Over Threefold

**DOI:** 10.64898/2026.01.31.702778

**Authors:** Roman Teo Oliynyk, George M. Church

## Abstract

We demonstrate DNA circularization efficiencies more than threefold greater than those predicted by classical Jacobson–Stockmayer theory for the ligation of linear double-stranded DNA. To quantify deviations from classical expectations, we experimentally calibrated the J-factor by ligating 452 bp DNA fragments bearing pre-cut, purified 4-nt overhangs at the optimal shortest minicircle length and at the highest DNA concentration; this reference value then enabled accurate calculation of expected cyclization efficiencies across the full range of DNA lengths (452–952 bp) and the three concentrations examined. These elevated cyclization efficiencies are enabled by simultaneous restriction cutting and ligation using the Type IIS enzyme BsaI-HFv2 in combination with T4 DNA ligase under finely optimized buffer conditions, yielding a 3.4-fold improvement over the classical estimate and achieving 75% cyclization at a high, practically relevant DNA concentration of 120 ng/*µ*l. We also identified a second enzyme, Esp3I, that exceeds the classical expectations by 2.3-fold (though less efficiently than BsaI-HFv2), while BbsI cut-ligation systematically underperformed expectations, providing insight into possible mechanisms underlying the outstanding performance of the first two enzymes. These results reveal a biologically mediated exception that overcomes the long-standing mechanistic expectations of Jacobson– Stockmayer theory and highlight the value of systematically screening enzyme combinations to discover additional systems capable of highly efficient intramolecular ligation.

## 1. Introduction

Circular DNA has emerged as a widely used platform for delivering genome-editing constructs [1] owing to its markedly greater stability in the cytoplasm compared with linear DNA [2]. Plasmid DNA was among the earliest delivery vehicles for genome editing [3–5]; however, plasmids include a bacterial backbone that increases construct size and can produce detrimental effects in sensitive cell types [6]. Minicircles – circular DNA produced without a bacterial backbone – offer improved transfection efficiency, enhanced nuclear transport, and higher transcription levels relative to conventional plasmids [7,8]. Renewed interest in minicircles over the past few years, exemplified by the 2024 study “Minicircle DNA vectors: A breakthrough in non-viral delivery of CRISPR base editors?” [9] and related work [10–12], underscores the growing recognition of their potential and the pressing need for efficient, scalable production methods.

Small minicircles are particularly attractive for delivering individual guide RNA (gRNA) expression cassettes for CRISPR–Cas9, which are usually ∼450 bp in size; we have published such a method [13,14] for applications in traditional CRISPR–Cas9 [15] as well as for base and prime editing [16–18]. Previously [13], we experimentally demonstrated a circularization protocol that achieved a yield of 62% (even with spin-column cleanup losses) for ∼450 bp minicircles at a high input DNA concentration of 120 ng/*µ*l, exceeding expectations set by Jacobson–Stockmayer theory [19–21]. In the present study we investigate the factors and mechanisms that may underlie these enhanced efficiencies.

Since the seminal Jacobson–Stockmayer formulation of the J-factor in 1950 [19], numerous theoretical and experimental studies have examined DNA circularization and polymerization kinetics [22–28]. The basic principle states that, at sufficient DNA length and concentration, linear polymers predominate, whereas low concentration and short chain length favor intramolecular cyclization. Notably, exceptions to the classical theory have been reported in polymer chemistry – for example, critiques and alternative syntheses of cyclic polymers that highlight limitations of the Jacobson–Stockmayer framework [29].

Calculations based on the J-factor predict far lower circularization efficiencies than the high yields we originally reported using a simultaneous cut-and-ligation method [13]. There we found that the specific combination of the Type IIS restriction enzyme BsaI-HFv2 with T4 DNA ligase in a simultaneous cut-and-ligation reaction yields exceptionally high circularization efficiency. We hypothesize that this dramatic enhancement arises from the ability of certain Type IIS restriction enzymes to simultaneously engage both recognition sites on the same DNA molecule before releasing the DNA (a process known as *cis* synapsis) [30]. This behavior leads to transient colocalization (processive in *cis*) of the two newly generated DNA ends, dramatically increasing their local effective concentration and thereby facilitating efficient intramolecular ligation by T4 DNA ligase – far beyond what random diffusional encounters between completely free DNA ends would allow under classical Jacobson–Stockmayer assumptions. In this study we empirically determined the J-factor corresponding to the ligation of linear double-stranded DNA (dsDNA) with pre-cut overhangs at the optimal minicircle size of 452 bp and compared our method performance with two additional Type IIS restriction enzymes, BbsI and Esp3I, thereby providing insight into possible mechanisms that lead to outstanding cyclization performance.

We present a further optimized simultaneous cut-and-ligation method that improves upon prior performance within the previously reported practical length range (452–952 bp). At the optimal minicircle size of 452 bp these refinements yield circularization efficiencies 3.4 times higher than the cyclization efficiency measured using pre-cut and purified dsDNA overhangs (which is, equivalently, the maximum estimate consistent with the Jacobson– Stockmayer model), even at high DNA concentrations suitable for practical applications.

These results demonstrate that the enzyme pair BsaI-HFv2 (and to a lesser extent Esp3I) together with T4 DNA ligase, under optimized buffer conditions, constitutes a biologically derived exception that exceeds the long-assumed constraints on DNA cyclization efficiency.

## 2. Materials and Methods

### 2.1. Overview and Optimizations to the Original Protocol

The principle and steps of the simultaneous cut-ligation method are schematically presented in Figure 1; for specifics, refer to the figure legend and the full protocol [14]. Experiments used the identical dsDNA sequences as in the original report [13], spanning the 452–952 bp range, with a 12-hour circularization reaction performed at a constant 37°C to minimize mismatch ligation [31,32]. The ATP concentration was set to 2 mM (within the optimum range for T4 DNA ligase) [33], which we experimentally demonstrated [13] sustains ligation activity throughout the 12 h reaction; repeated restriction-enzyme cutting and re-ligation events could otherwise deplete available ATP and limit the formation of circular concatemers.

**Figure 1.**
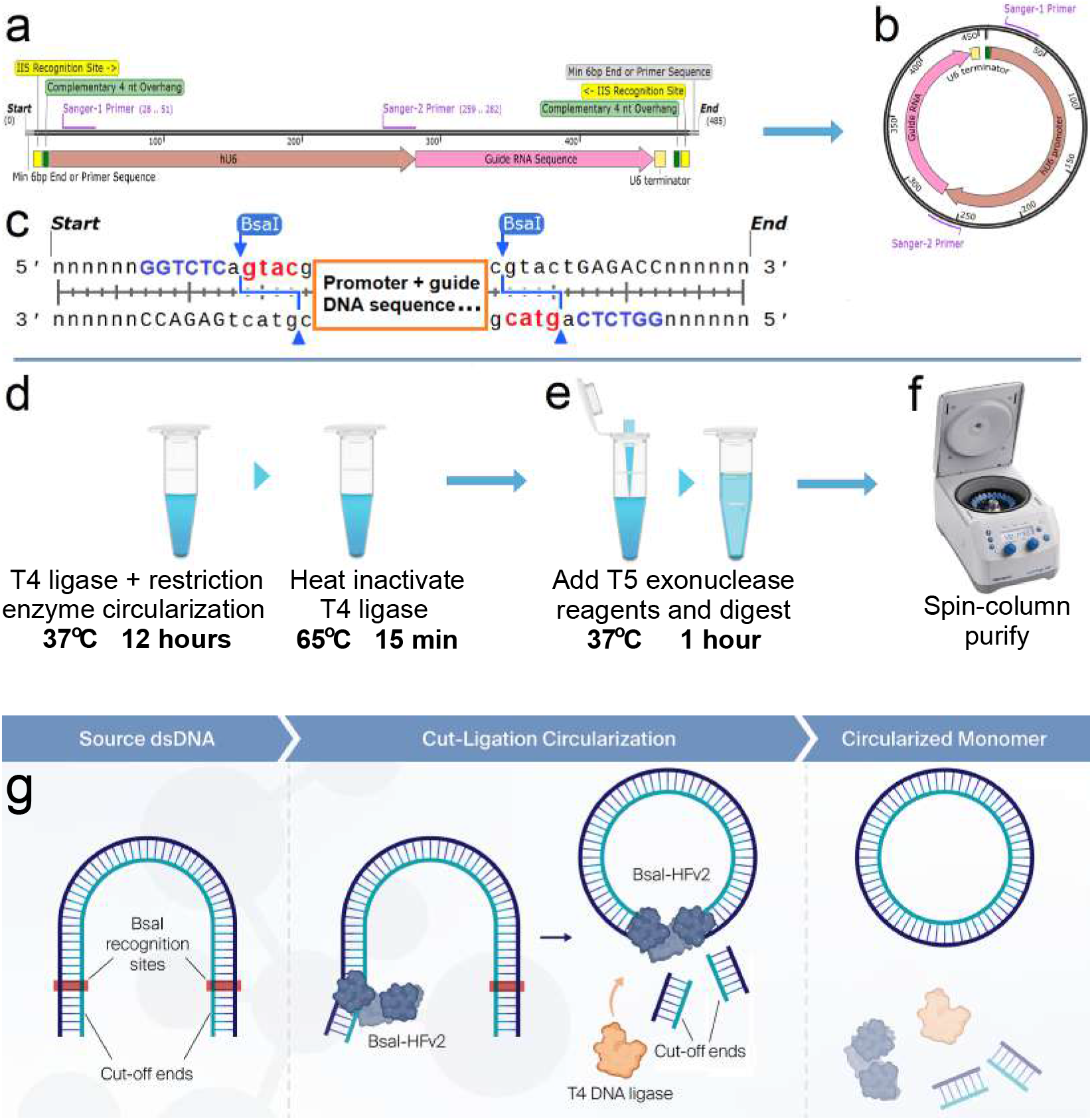
Simultaneous cut-ligation method and circularization steps. (**a**) dsDNA containing type IIS restriction enzyme recognition sites with complementary overhangs on both ends of the DNA fragment. (**b**) Cuts by the restriction enzyme and ligation of the resulting overhangs form the circular DNA expression element. (**c**) Schematic view of complementary 4nt overhangs. *Method steps:* (**d**) Circularization. (**e**) T5 exonuclease digestion of non-circular DNA in one-pot reaction. (**f**) Spin column cleanup. (adapted from Oliynyk & Church (2022)[13]) (**g**) Proposed mechanism of enhanced circularization efficiency in the simultaneous cut-ligation reaction employing BsaI-HFv2; the comparative performance and mechanistic details are explored in the Discussion.

To validate higher circularization efficiency at lower DNA concentrations, in addition to the standard concentration of 120 ng/*µ*l, we also performed experiments at 60 ng/*µ*l and 30 ng/*µ*l – concentrations expected to exhibit higher circularization efficiency. We followed the published conditions, with slight but important modifications to reagent formulations in the T5 exonuclease digestion step that further improved recovery of circular products compared to previously published yields [13] for the smallest minicircle sizes (452 and 552 bp).

### 2.2. Experimental

#### 2.2.1. Fine-tuning the Digestion of Linear DNA

The previously published methods [13,14] were designed to provide a robust, reliable, and user-friendly approach for producing circular DNA vectors, prioritizing simplicity and consistency for routine laboratory applications.

In the present study, our primary objective was to explore and demonstrate the maximum achievable circularization efficiency under optimized conditions. To this end, we introduced targeted refinements to the reaction buffers, aimed at minimizing potential over-digestion by T5 exonuclease – particularly for the smallest, highly curved minicircles that may exhibit transient structural breathing due to A/T-rich regions and potential hairpin-forming sequences in the hU6 promoter [34,35]. These modifications include reversion to the NEB T4 DNA ligase buffer (pH 7.5) for the circularization step and supplementation of the digestion buffer with 60 mM potassium acetate (KOAc) to enhance DNA stability, reduce breathing, and moderately attenuate T5 exonuclease activity while preserving efficient removal of linear DNA.

The modified buffer compositions are summarized below (Tables 1 and 2). For comprehensive step-by-step instructions, including thermocycling parameters, reagent preparation, and general procedures, readers are referred to the original published method [13,14].

**Table 1.** Circularization reaction ingredients.

| Component | 100 μl reaction | Final conc. |
| --- | --- | --- |
| dsDNA - 300 ng/μl | 10, 20, 40 μl | 30, 60, 120 ng/μl |
| 10X T4 DNA Ligase Buffer (10X B0202S) | 10 μl | 1X |
| T4 DNA Ligase (NEB M0202L, 400 units/μl) | 6 μl | 24 units/μl |
| BsaI-HFv2 (NEB R3733L, 20 units/μl) | 6 μl | 1.2 units/μl |
| ATP (NEB P0756S, 10 mM) | 10 μl | 1 mM |
| Nuclease-free $H_2O$ (Invitrogen 10977015) | up to 100 μl | n/a |

**Table 2.** Digestion reaction ingredients to add to each 100 *µ*l of circularization reaction after heat inactivation for digestion step (Figure 1e)

| Component | Per 100 $\mu\text{l}$ reaction | Final conc. |
| --- | --- | --- |
| **30 ng/$\mu\text{l}$ input DNA** |  |  |
| KOAc 1M (Thermo Fisher AAJ62817AE, pH 7.5) | 6.5 $\mu\text{l}$ | 60 mM |
| T5 Exonuclease (NEB M0663, 10 units/$\mu\text{l}$) | 2 $\mu\text{l}$ | 0.2 units/$\mu\text{l}$ |
| **60 ng/$\mu\text{l}$ input DNA** |  |  |
| KOAc 1M (Thermo Fisher AAJ62817AE, pH 7.5) | 6.6 $\mu\text{l}$ | 60 mM |
| T5 Exonuclease (NEB M0663, 10 units/$\mu\text{l}$) | 4 $\mu\text{l}$ | 0.4 units/$\mu\text{l}$ |
| **120 ng/$\mu\text{l}$ input DNA** |  |  |
| KOAc 1M (Thermo Fisher AAJ62817AE, pH 7.5) | 12.0 $\mu\text{l}$ | 60 mM |
| T5 Exonuclease (NEB M0663, 10 units/$\mu\text{l}$) | 8 $\mu\text{l}$ | 0.4 units/$\mu\text{l}$ |
| Tris 1M (Invitrogen 15567027, pH 7.5) | 2.5 $\mu\text{l}$ | 25 mM |
| MgCl<sub>2</sub> 1M (Technova M0303) | 1.0 $\mu\text{l}$ | 10 mM |
| Nuclease-free H<sub>2</sub>O (Invitrogen 10977015) | up to 100 $\mu\text{l}$ | n/a |

If the buffer is prepared for larger volumes or multiple digestion reactions, these ingredients may be scaled and combined, and added in specified combined volume to circularization reaction. For reactions with input DNA concentrations of 120 ng/ $\mu\text{l}$ , the reaction mixture is added in equal quantity to the ligation reaction buffer, thus KOAc and T5 exonuclease are calculated for 2X volume, while Tris and MgCl<sub>2</sub> compensate for the dilution of these components in the ligation reaction.

Through experimental fine-tuning of the digestion buffer components and T5 exonuclease concentrations, we achieved complete digestion of all non-circular DNA. Digestion time-course gels revealed traces of high-molecular-weight concatemers remaining in some samples after 30 min, with complete removal of linear species by 60 min and no further changes in band patterns after 90 min (Figure 2a–d), confirming that 60 min is sufficient for exhaustive digestion of non-circular DNA for all concentration-optimized digestion buffer compositions.

**Figure 2.**
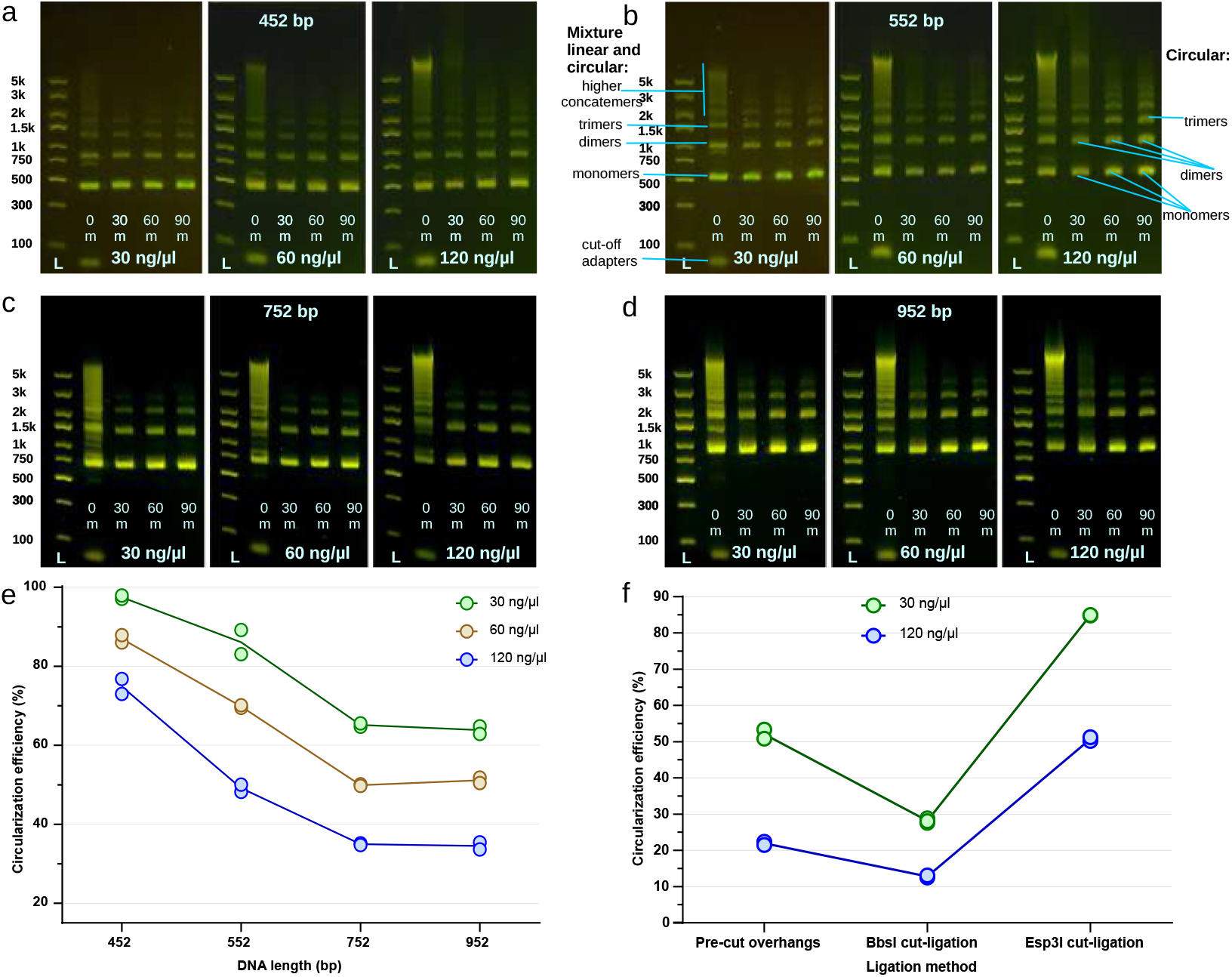
Circularization efficiency and DNA band distribution following circularization and validation of T5 exonuclease digestion. (**a–d**) Agarose gels showing the time-course of T5 exonuclease digestion for the four minicircle lengths (452, 552, 752, 952 bp) each containing subplots at 30, 60 and 120 ng/*µ*l concentrations of source dsDNA (unmodified gel photos are available in Supplementary Data). “L” = gel ladders. “0 min” lanes = product immediately after circularization step (heat-inactivated, no T5 added yet). Subsequent lanes = 30 min, 60 min and 90 min of T5 digestion (each sample purified on spin column before loading). See Supplementary Data for unmodified gel images. (**e**) DNA circularization efficiency for cut-ligation with BsaI-HFv2 + T4 DNA ligase. (**f**) DNA circularization efficiency for DNA with pre-cut overhangs, and cutting using BbsI and Esp3I. (In (**e**) and (**f**) corrected for column recovery losses; n=2 duplicate reactions, individual values shown).

Here we provide a summary of the ingredients that were modified for better performance; refer to [14] for detailed instructions. Sufficient dsDNA was prepared by PCR as described in our 2025 method [14] and DNA concentration adjusted to 300 ng/*µ*l. This DNA was combined into the circularization buffer in Table 1 in quantities 10 *µ*l for samples with concentration 30 ng/*µ*l, 20 *µ*l for 60 ng/*µ*l and 40 *µ*l for 120 ng/*µ*l, correspondingly. The temperatures and thermocycler timings are specified in Figure 1d,e. After completion of the circularization step (Figure 1d), the Table 2 ingredients are added to the one-pot reaction tube for 30 ng/*µ*l and 60 ng/*µ*l input dsDNA concentrations, proceeding per Figure 1e, followed by spin-column cleanup Figure 1f using QIAquick PCR Purification Kit (QIAGEN 28106).

As a part of finetuning, we found that with the intended lower T5 exonuclease activity, digestion of reactions containing 120 ng/*µ*l input dsDNA was not complete within one hour, likely owing to the high concentration of short linear fragments – byproduct of digestion, though full digestion was still achieved if digestion time was extended. To address this, we doubled the effective reaction volume for these samples by adding an equal amount of the digestion mix formulated in the third section of Table 2. This adjustment fully resolved the issue, achieving complete digestion within the intended one-hour timeframe.

These buffer optimizations successfully mitigated the risk of over-digestion highlighted above. Compared to the original method [13], the refined digestion step improved recovered yields for the shortest minicircles at high input DNA, while yields for longer minicircles remained essentially unchanged (see in Supplementary Data), confirming that the moderated exonuclease activity and enhanced DNA stability effectively protected shorter minicircle products from gradual degradation. Another indication of the well-adjusted digestion method is virtually no DNA loss for the shortest minicircles at the lowest concentration; the gel image showed virtually complete circularization with minimal shadow of higher-order concatemers (Figure 2a, lane 0 min), matching the spin-column-recovered yield after digestion step and accounting for the experimentally determined column recovery rate. Taken together, these results validate the buffer adjustments as both protective against over-digestion of circular products and permissive for complete elimination of linear species.

The Excel spreadsheets for calculating and scaling reaction ingredients are available in the Supplementary Data.

#### 2.2.2. Cut-ligation using other type IIS restriction enzymes

The BbsI-HF (NEB #R3539L) and Esp3I (NEB #R0734L) type IIS restriction enzymes were used as described in the above section, substituting them in identical number of activity units for BsaI-HFv2 (see the dsDNA designs in Supplementary Data *452bp-BbsI*.*gbk* and *452bp-Esp3I*.*gbk*).

#### 2.2.3. Preparation and circularizing of dsDNA with pre-cut overhangs

Ligating the 452 bp minicircles with 4-nt single-stranded overhangs identical to our cut-ligation protocol provided the empirical value of cyclization efficiency and allowed determination of the corresponding Jacobson–Stockmayer J-factor that by definition characterizes it.

To simplify the preparation of such DNA we designed an additional dsDNA with longer flanking cut-off arms of 100 and 150 bp, which allowed for the ease of determination of the completeness of cutting the overhangs and for gel purification of the 452 bp band while eliminating the cut-offs (see file *452bp-With-Padded-CutOffs*.*gbk* in Supplementary Data).

The preparation of pure dsDNA fragments with overhangs for production of 452 bp minicircles was performed following standard molecular techniques and the protocol described in the main article:

- Restriction enzyme cutting of the source DNA containing two BsaI recognition sites using NEB BsaI-HFv2 enzyme and rCutSmart buffer, validating the completeness of digestion using Invitrogen E-Gels and purification using QIAGEN QIAquick spin column kit.
- Gel purification using Owl EasyCast B2 Mini Gel Electrophoresis Systems followed by the QIAGEN QIAquick gel extraction spin column kit.
- Second round of QIAGEN QIAquick spin column kit purification to eliminate residual contaminants from the above gel extraction, resulting in pure DNA with 4-nt matching overhangs.
- Performing a 12-hour ligation using the protocol described in the Methods in the main article, substituting the BsaI-HFv2 by an equal amount of water.

### 2.3. Theoretical: Jacobson–Stockmayer Estimates of DNA Circularization Efficiencies

Circularization efficiencies for linear double-stranded DNA fragments of 452, 552, 752, and 952 bp were estimated theoretically using the worm-like chain (WLC) model [20] and Jacobson-Stockmayer (JS) theory [19].

The Jacobson-Stockmayer factor (*J*), defined as the effective local concentration of one DNA end in the vicinity of the other (in molar units) [19], was approximated in the Gaussian (long-chain) limit of the WLC model [20]. Per Yamakawa (2016) [21] equation 7.39, the *J* -factor for longer dsDNA lengths *L* can be approximated by

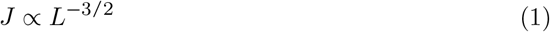

with *L* the DNA contour length (proportional to base-pair number). This power-law scaling arises from the probability density of end-to-end closure in three-dimensional random-walk chains in the large-*N* limit [19,21].

The numerical computations are based on ring-closure probabilities in the twisted WLC model applied specifically to dsDNA parameters (persistence length *P* ≈ 50 nm or ∼147 bp, helical repeat ∼10.5 bp/turn) [20]. The Shimada–Yamakawa calculations [20] show a broad peak in the effective concentration (directly related to the *J* -factor) in the range of tens of nM for contour lengths corresponding to ∼400–600 bp (for illustration see Fig. 5 in Shore et al. [23]), with the Gaussian *L*^−3/2^ approximation providing a good fit in this regime and beyond for lengths ≫ 2*P*.

From our experimental results, circularization efficiency is highest for the 452 bp fragment. To calibrate the J-factor at this optimal length, we used the empirical circularization efficiency of 21.9% (accounting for all concatemers) obtained with pre-cut and purified overhangs at a DNA concentration of 105.5 ng/*µ*l (see Table 4). Using the *Efficiency from J-Factor* sheet in the Supplementary Data spreadsheet *Circularization Efficiency*.*xlsx*, we entered the known DNA molar concentration from Table 4 and iteratively adjusted the J-factor value until the calculated circularization efficiency matched the observed 21.9%. This calibration yields a J-factor of 86.0 nM. Thus, for all other lengths, *J* -factors were normalized relative to this maximum value at the shortest fragment (*L*_0_ = 452 bp).

For fragments of different lengths *L*_*n*_, the relative scaling factor is *x* = *L*_0_/*L*_*n*_ (where *L*_0_ = 452 bp is the optimal shortest length). The J-factor is then

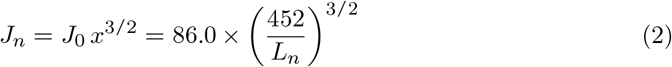

(in nM); the calculated *J* -factors (in nM) used here are presented in Table 3.

**Table 3.** Calculated *J* -factors per DNA length.

| DNA Length (bp) | $J$-factor (nM) |
| --- | --- |
| 452 | 86.0 |
| 552 | 63.7 |
| 752 | 40.1 |
| 952 | 28.1 |

Because we started with the experimentally determined circularization efficiency for 452 bp minicircles, Equation 2 provides the relative J-factor change; when entered into Equation 3 and Equation 4, it yields the relative change in circularization efficiency for the set of DNA lengths and concentrations shown in Table 5 and Table 6. This ensures that the circularization efficiency estimates remain invariant under differing WLS model parameters.

DNA mass concentrations (ng/*µ*L) for each fragment length and experimental condition are presented in Table 4.

**Table 4.** DNA mass concentrations (ng/*µ*L) and molar concentrations *C* for each fragment length.

| DNA lengths (bp) | DNA concentration |  |  |
| --- | --- | --- | --- |
|  | Low | Medium | High |
| Mass concentration of circularizable DNA (ng/μL) | | | |
| 452 | 26.4 | 52.8 | 105.5 |
| 552 | 27.0 | 53.9 | 107.9 |
| 752 | 27.7 | 54.4 | 110.9 |
| 952 | 28.2 | 56.3 | 112.7 |
| Molar concentration of circularizable DNA (nM) |  |  |  |
| 452 | 88.5 | 177.0 | 353.6 |
| 552 | 74.1 | 147.9 | 296.2 |
| 752 | 55.8 | 109.6 | 223.4 |
| 952 | 44.9 | 89.6 | 179.4 |

The input DNA concentrations are 30, 60 and 120 ng/ $\mu$ L, however as the input DNA is 62 bps longer to account for restriction enzyme sites and PCR adapters, which are cut off during the ligation/overhang cutting step. The resulting concentrations are smaller and slightly vary for each condition, thus for clarity here and later we call them low, medium and high concentrations. Corresponding molar concentrations $C$ (nM), calculated using an average dsDNA molar mass of 660 g/mol per base pair.

The percentage efficiency for formation of monomeric circles was calculated from the kinetic competition between intramolecular cyclization (first-order rate proportional to *J*) and intermolecular dimerization (second-order rate proportional to molar concentration *C*), yielding the circular monomer fraction [19,36]:

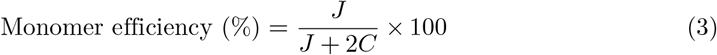

The factor of 2 accounts for the stoichiometry of dimer formation consuming two molecules per ligation event.

For circular concatemers formed from an integer number *m* ≥ 2 of identical monomer fragments (total contour length *mL*_*n*_), the *J* -factor scales as

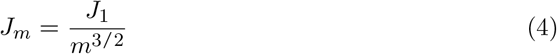

according to the Gaussian-regime prediction of JS theory [19], applied in the same manner as in Equation 1 and Equation 2.

Yields of circular concatemers (dimers, trimers, and higher-order species up to 16-mers) were estimated sequentially using the same kinetic branching model. See estimated efficiencies for a circular monomer in Table 5 and total circularization efficiency combining monomers and all circular 2–16-mers in Table 6 (see the *Efficiency from J-Factor* sheet in the Supplementary Data spreadsheet *Circularization Efficiency*.*xlsx*).

**Table 5.** Monomeric circularization efficiency.

| DNA lengths (bp) | DNA concentration |  |  |
| --- | --- | --- | --- |
|  | Low (%) | Medium (%) | High (%) |
| 452 | 32.7 | 19.5 | 10.8 |
| 552 | 30.1 | 17.7 | 9.7 |
| 752 | 26.4 | 15.5 | 8.2 |
| 952 | 23.8 | 13.6 | 7.3 |

Calculations for fraction of circular dimers and trimers are also presented in Supplementary Data spreadsheet.

These values represent approximate theoretical yields for the specified circular species in an irreversible ligation reaction; the remainder consists of linear oligomers (which are subsequently eliminated by T5 exonuclease digestion and spin-column purification steps). All percentages reflect the proportion of original DNA mass (or equivalently, incorporated original segments) in each circular form. Total circular efficiency (Table 6) is the sum of monomeric circles and all circular concatemers (higher-order species beyond 16-mers contribute negligibly).

**Table 6.** Total circularization efficiency as a sum of monomers and all circular 2–16-mers, used for comparison with experimental results in Table 8.

| DNA lengths (bp) | DNA concentration |  |  |
| --- | --- | --- | --- |
|  | Low (%) | Medium (%) | High (%) |
| 452 | 59.8 | 38.1 | 21.9 |
| 552 | 55.7 | 34.8 | 19.7 |
| 752 | 49.9 | 30.7 | 16.8 |
| 952 | 45.6 | 27.1 | 14.9 |

## 3. Results

### 3.1. Improved Experimental Results with Simultaneous Cut-Ligation With BsaI-HFv2 + T4 DNA Ligase

The modified digestion buffer efficiently eliminates linear DNA, as seen in the ligation/digestion test for all DNA lengths and three DNA concentration scenarios (30, 60 and 120 ng/*µ*l). For this test, we performed the ligation using reagents listed in Materials and Methods Table 1 followed by heat inactivation (Figure 1d), shown as digestion time 0 min – no digestion – on the plots Figure 2a–d. As per Jacobson-Stockmayer theory [37], for the DNA with matching overhangs, at lower concentrations circularization will be favored, as diffusional collision in cis is more likely than in trans, however some fraction of concatemers is always likely. At higher concentrations, the likelihood of trans collision of matching overhangs and their ligation increases. It is particularly notable that when linear dimers are produced, they are even less likely to circularize rather than ligate further in trans, and polymerization probability progresses even more for trimers and higher concatemers, thus accumulating higher linear concatemers observed above 5 kbp, as seen in Figure 2a–d at 0 min lanes.

The 0 min digestion (before digestion) lanes in Figure 2a–d show that at 452 bp the circularization is most effective, resulting in the smallest amount of linear high-molecular-weight concatemers compared to the three longer DNA. There is only the faintest trace of linear product for 452 bp minicircles after the ligation step – lane 0 min in Figure 2a at DNA concentration of 30 ng/*µ*l – indicating almost 100% circularization efficiency; at 60 ng/*µ*l the linear high-molecular-weight concatemers product in lane 0 min becomes more prominent, and even more so at 120 ng/*µ*l. The same observation applies to the circularization efficiency of longer minicircles, where the amount of linear concatemers in the 0 min bands increases with minicircle size even for the lowest input DNA concentration (Figure 2b–d).

Then, we proceeded to the digestion step (Figure 1e), adding T5 Exonuclease and associated reagents as per Materials and Methods Table 2, and taking samples for gel imaging at 30 min intervals, each sample spin-column purified (Figure 1f) before gel imaging. There is almost complete disappearance of the high-molecular-weight linear concatemers after 30 min digestion, with remaining non-circular DNA completely digested by 60 min, and the band pattern remaining unchanged for 90 min digestion samples, indicating that 60 minutes of T5 exonuclease digestion formulations are adequate for the purposes of our method.

The simultaneous cut-ligation method was performed in duplicate for all 4 DNA lengths and three concentrations, with the results reported in Figure 2e and Table 8. Our validation experiments of the QIAquick spin-column efficiency using the 452 bp minicircle product containing all circular concatemers as input showed DNA recovery after one cycle of cleanup equal to 0.88, which we rounded to 0.9 (this matches the QIAGEN manual estimates). Here we are primarily interested in pure circularization efficiency, thus the yields are adjusted by this value to compensate for the cleanup losses (essentially, dividing the measured yield of the DNA product by 0.9). Higher circularization efficiency is observed near the optimum J-factor for DNA sizes of ∼452 bp (see also explanation in Materials and Methods); thus, at DNA concentration 30 ng/*µ*l circularization efficiency is the highest (97.5%) at 452 bp and lower for each longer DNA length (see Table 8). At DNA concentration 60 ng/*µ*l the circularization efficiency is lower than for 30 ng/*µ*l, and lower yet at 120 ng/*µ*l, still remaining impressively high for all four DNA sizes (seen in Figure 2e and Table 8). The lower the circularization efficiency reported in Figure 2e and Table 8, the higher the visible amount of high-order concatemers seen in the corresponding 0 min lanes (post-circularization but pre-digestion) in Figure 2a–d.

### 3.2. Demonstration of Complete Digestion of Unreacted Linear DNA by T5 Exonuclease Digestion

While the T5 exonuclease digestion results above showed complete removal of high-molecular-weight linear concatemers, an additional control experiment presented in Figure 3a– b demonstrates that even if no circularization occurred, any remaining linear DNA would still be fully digested during the T5 exonuclease step. We performed a mock reaction using a linear 514 bp dsDNA construct that lacks both restriction enzyme recognition sites and matching overhangs (*452bp-Esp3I*.*gbk*; see Supplementary Data) at 120 ng/*µ*l, thereby ensuring the DNA would remain linear after the reaction. In Figure 3a we tested the ligation buffer alone (BsaI-HFv2 replaced by an equal volume of water). In Figure 3b we used the standard cut-ligation buffer containing BsaI-HFv2. Lane 1 in both panels shows the original linear dsDNA as a single band, as expected. The reactions were incubated at 37°C for 12 h, followed by heat inactivation at 65°C for 15 min. Because the dsDNA lacks matched overhangs (and the Esp3I sites in panel b are not recognized by BsaI-HFv2), the DNA remains linear in both cases. Each sample was then split into two equal aliquots. One aliquot was purified directly by spin column (lane 2). The second aliquot was subjected to 1 h of T5 exonuclease digestion per our protocol and then purified (lane 3). Lanes 2 in Figure 3a–b show no additional bands, confirming that no circularization occurred. Lanes 3 are empty, even though four times the volume of digestion reaction product was loaded compared to lanes 2, demonstrating that the T5 exonuclease digestion step completely eliminates any unreacted linear DNA. Notably, when circularization efficiency is high (as in Figure 2a), the mass of linear DNA requiring digestion is minimal, making the digestion step even more rapid and efficient.

**Figure 3.**
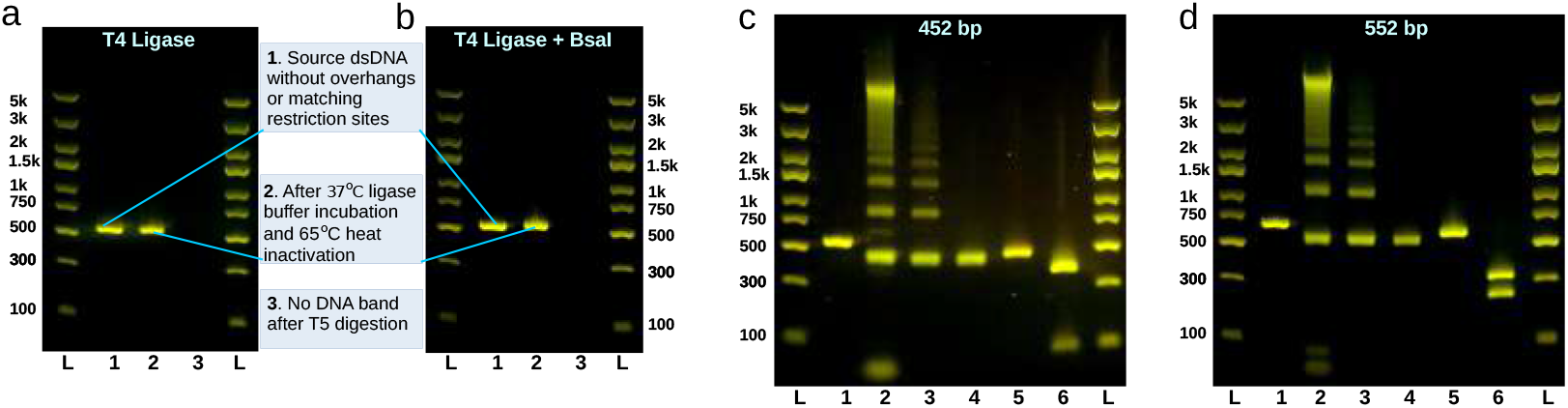
T5 digestion and circularization validation experiments. **(a)–(b)** Complete digestion of unreacted linear DNA by T5 exonuclease: (**a**) Mock reaction of non-circularizable DNA without BsaI-HFv2. (**b**) Mock reaction of non-circularizable DNA using the standard cut-ligation buffer. Lane 1: original linear dsDNA. Lane 2: product after incubation and heat inactivation (no circularization expected). Lane 3: product after T5 exonuclease digestion (no DNA band remains, confirming complete digestion of linear DNA). “L” - gel ladders. **(c)–(d)** Restriction enzyme validation of circular monomer purity for 452 bp (**c**) and 552 bp (**d**) minicircles. Lane 1: original linear dsDNA. Lane 2: circularization product before T5 digestion. Lane 3: product after T5 digestion. Lane 4: gel-purified circular monomer band. Lane 5: single-cut digestion of monomer with NdeI (single linear band, confirming circularity). Lane 6: double-cut digestion (expected fragment sizes: 86 bp + 366 bp for 452 bp (two sites for AerI); 237 bp + 315 bp for 552 bp (NdeI and ApoI).

### 3.3. Demonstration of Circular Monomer Purity by Restriction Enzyme Cutting

In our original publication we presented two PCR-based assays to validate circularization. Here we show an alternative approach using restriction enzyme digestion on gel-purified circular monomer bands for the 452 bp and 552 bp minicircles (Figure 3c–d). Lane 1 in each panel shows the corresponding initial linear dsDNA as a single band. Lane 2 shows the circularization product before T5 digestion. Lane 3 shows the same material after T5 exonuclease digestion, which has eliminated all linear concatemers and unreacted linear DNA. Lane 4 shows the gel-extracted circular monomer band, which migrates at the expected position and matches the band in lane 3.

In lanes 5 we performed a single-cut digestion of the purified monomer using NdeI (NEB R0111S), which cuts approximately in the middle of the original dsDNA sequence (see *452bp-Circularized*.*gbk* and *552bp-Circularized*.*gbk* in Supplementary Data). If circularization had failed or been incomplete, multiple bands would appear; instead, both lanes 5 show a single linear band that migrates slightly slower than the circular band in lane 4, confirming that the product is fully circular.

In lanes 6 we performed double-cut digestions for further verification. For the 452 bp circle (Figure 3c) we used AerI (NEB R0528S), which has two recognition sites and produces fragments of 86 bp and 366 bp, exactly as observed. For the 552 bp circle (Figure 3d) we used NdeI and ApoI-HF (NEB R3566S), each with one recognition site, yielding the expected fragments of 237 bp and 315 bp, again matching the observed bands.

### 3.4. Comparison of Circularization Efficiency of DNA With Pre-cut and Purified Overhangs

By definition, the ligation efficiency of double-stranded DNA (dsDNA) with matching overhangs is characterized by the Jacobson–Stockmayer J-factor. We experimentally determined the circularization efficiency using DNA with pre-cut and purified overhangs designed to produce 452 bp circular vectors identical to our cut-ligation scenarios (see Table 7 and Figure 2f). The circularization efficiency was found to be 21.9% at a DNA concentration of 105.5 ng/*µ*l (which is equivalent to 120 ng/*µ*l DNA used for the cut-ligation reaction per Table 4). Based on the calculations presented in Materials and Methods, the J-factor corresponding to such circularization rate was determined to be 86.0 nM. Knowing this empirical J-factor, we can now apply it to calculating the expected circularization rates for all lengths and DNA concentrations tested, as it represents the upper limit of what Jacobson-Stockmayer theory would expect.

**Table 7.**
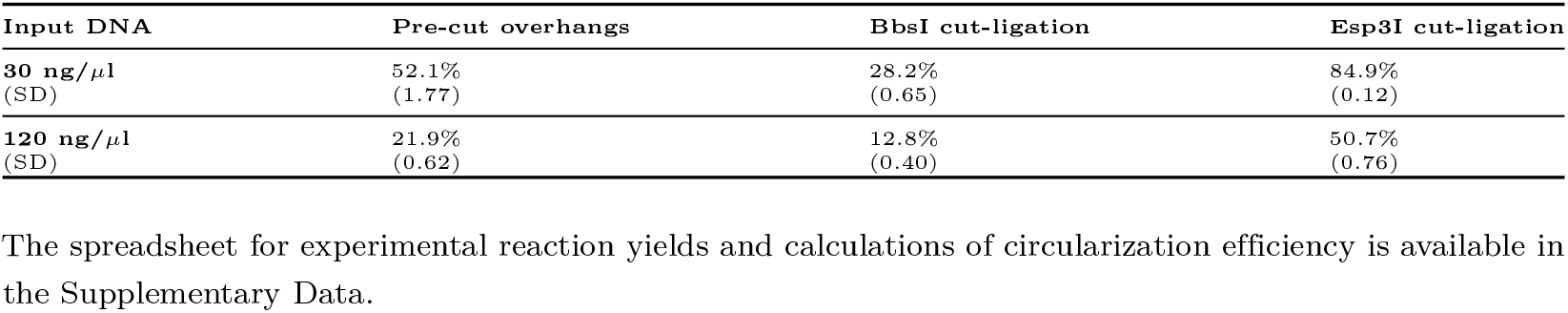
Experimental circularization efficiency of the ligation of the pre-cut overhangs and cut-ligation using BbsI-HF and Esp3I enzymes.

### 3.5. Circularization Efficiency of Simultaneous Cut-Ligation With BbsI-HF and Esp3I

In marked contrast to the above results, reactions under the same conditions with BbsI-HF (NEB #R3539L) yielded lower circularization efficiencies than ligation of DNA with pre-cut overhangs (see Table 7). At a DNA concentration of 120 ng/*µ*l, BbsI cut-ligation yielded only 12.8% minicircle DNA – approximately 42% lower than the 21.9% obtained with pre-cut overhangs, and more than fivefold lower than with BsaI-HFv2. This reveals that simultaneous cut-ligation with some enzymes can reduce circularization yield relative to pre-cut controls, in contrast to the higher yield observed with BsaI-HFv2. This also translates to an approximately 5-fold lower cyclization efficiency when using the BbsI-HF restriction enzyme compared to BsaI-HFv2 at a DNA concentration of 120 ng/*µ*l (as we noted earlier [14]).

A new positive discovery is Esp3I (NEB #R0734L), which produced circularization efficiencies substantially higher than those achieved by ligation of DNA with pre-cut overhangs and the corresponding J-factor, showing 2.3-fold higher circularization at 120 ng/*µ*l (see Table 7 and Figure 2f). While this efficiency is lower than that of BsaI-HFv2, taken together with the lower performance of BbsI-HF, these results motivate a hypothesis for the mechanism underlying the enhanced performance of our protocol, which we present in the Discussion.

### 3.6. Summary Table of the Experimental Results with Simultaneous Cut-Ligation With BsaI-HFv2 + T4 DNA Ligase

Table 8 compares the experimental efficiency (first row in each concentration section) and the expected theoretical performance (second row in each concentration section). With this method, the combination of BsaI-HFv2 and T4 DNA ligase, when used in simultaneous cut-and-ligation with optimal buffer components, exceeds the physics-model circularization efficiency by 3.4-fold (75% vs 21.9% yield at the highest DNA concentration).

At the practical level, achieving high circularization at 120 ng/*µ*l for the shortest 452 bp minicircles makes this method by far the most efficient currently available for producing small circular DNA vectors, with common applications in gRNA expression for CRISPR-Cas genome editing [13,14].

**Table 8.** Experimental circularization efficiency of the optimized simultaneous cut-ligation method versus classical Jacobson–Stockmayer physics predictions. The BsaI-HFv2 + T4 DNA ligase combination exceeds theoretical expectations 3.4-fold at high input DNA concentrations for the shortest minicircles.

| Input DNA | Data source | Circularization efficiency per minicircle size |  |  |  |
| --- | --- | --- | --- | --- | --- |
|  |  | 452 bp | 552 bp | 752 bp | 952 bp |
| 30 ng/μl | Method | 97.5% | 86% | 65% | 64% |
|  | Model estimate | 59.8% | 55.7% | 49.9% | 45.6% |
| 60 ng/μl | Method | 87% | 70% | 50% | 51% |
|  | Model estimate | 38.1% | 34.8% | 30.7% | 27.1% |
| 120 ng/μl | Method | 75% | 49% | 35% | 34% |
|  | Model estimate | 21.9% | 19.7% | 16.8% | 14.9% |

## 4. Discussion

The primary conclusion of this research is that, after more than 75 years during which the Jacobson–Stockmayer J-factor has been regarded as the fundamental physical constraint on cyclization of flexible polymers, the specific combination of BsaI-HFv2 and T4 DNA ligase in a simultaneous cut-ligation reaction with optimized buffer conditions overcomes these presumed limitations for double-stranded DNA, delivering circularization efficiencies more than threefold higher than expected from classical theory.

The exceptional performance observed with the BsaI-HFv2 and T4 DNA ligase cut-ligation combination is further supported by the discovery of a second high-performing enzyme, Esp3I, which produced circularization efficiencies approximately 2.3-fold higher than those predicted from the J-factor based on ligation of DNA with pre-cut overhangs (although still lower than the 3.4-fold improvement observed with BsaI-HFv2 at 120 ng/*µ*l).

These results indicate that circularization outcome may depend on the dynamics of restriction-enzyme binding and cutting at two recognition sites on a single DNA molecule. Bath et al. (2002) [30] reported that Type IIS restriction enzymes such as BsaI and BsmBI, when two recognition sites are present on one molecule, often perform the second cut before releasing the position of the first cut. While NEB BsmBI was unsuitable for our simultaneous cut-ligation reaction because of its high optimal temperature (55°C) and incompatible buffer requirements, its isoschizomer Esp3I matched our reaction parameters and indeed showed outstanding performance. BbsI, which was not reported by Bath et al. to exhibit this property, underperformed relative to ligation of DNA with pre-cut overhangs by 42%.

We hypothesize that circularization efficiency improves when a restriction enzyme such as BsaI or Esp3I cuts both recognition sites before releasing the DNA, producing transient intra-segment colocalization (processive in cis) of the two DNA ends that facilitates ligation by T4 DNA ligase (see Figure 1g). The diminished results observed with BbsI suggest that this enzyme may cut one site at a time without promoting intra-segment colocalization, and may even transiently shield the second cut site, thereby favoring trans ligation and reducing circularization efficiency.

Reviewing Table 8, at our highest DNA concentration of 120 ng/*µ*l the BsaI + T4 ligase cut-ligation exceeds model estimates by 3.4-fold for 452 bp, 2.5-fold for 552 bp, 2.1-fold for 752 bp, and 2.3-fold for 952 bp. This shows that while the expected model efficiency diminishes with DNA length, the colocalization effect afforded by restriction enzyme cut-ligation may also diminish for longer DNA. The initial protocol was developed for the production of circular RNA expression vectors for CRISPR prime and base editing [13], with DNA circle sizes around 450 bp, and is therefore well suited to that application. As we demonstrated previously [13], the presence of a fraction of concatemers can also improve editing performance.

Therefore, the phenomenon we observe represents a biologically derived exception – and potentially a family of such exceptions – to the long-standing expectation of low DNA cyclization efficiency.

## 5. Conclusions

These findings demonstrate that the classical Jacobson–Stockmayer theory, while remarkably robust for more than seven decades, does not constitute an absolute physical barrier when specific enzyme systems are employed for DNA circularization. By showing that very high circularization efficiencies are achievable in practice, this work removes a long-standing conceptual constraint and should encourage the broader community to pursue systematic screening of restriction enzymes, ligases, and accessory factors to discover additional high-performance systems.

## Supplementary Materials

The supporting information can be downloaded as SupplementaryData.ZIP.

## Data Availability Statement

The unmodified gel image photos, the DNA sequences, the circularization efficiency spreadsheet and reaction reagents scaling spreadsheet are available in SupplementaryData.ZIP file.

## Statistics and Reproducibility

All protocol reactions were performed in independent reaction duplicates (n=2) as stated in figures, and yield percentage values were averaged.

## Acknowledgments

The authors express their sincere gratitude to Davide Michieletto (School of Physics and Astronomy, and MRC Human Genetics Unit, University of Edinburgh) for his thoughtful discussions and expert insights with the Jacobson–Stockmayer model calculations.

